# Oligodendrocyte-specific expression of *PSG8-AS1* suggests a role in myelination with prognostic value in oligodendroglioma

**DOI:** 10.1101/2023.12.06.570364

**Authors:** Maria de los Angeles Becerra Rodriguez, Elena Gonzalez Muñoz, Tom Moore

**Affiliations:** School of Biochemistry and Cell Biology, University College Cork, Cork, Ireland; SFI Centre for Research Training in Genomics Data Science, University College Cork, Cork, Ireland; Biomedical Research Institute and Nanomedicine Platform (IBIMA-BIONAND) 29590 Málaga, Spain; Department of Cell Biology, Genetics and Physiology, University of Malaga, 29071 Málaga, Spain

**Keywords:** oligodendrocyte, myelination, segmental duplication, human-specific gene, lncRNA, brain, glioma, pregnancy-specific glycoprotein, *PSG8-AS1*

## Abstract

The segmentally duplicated Pregnancy-specific glycoprotein (*PSG*) locus on chromosome 19q13 may be one of the most rapidly evolving in the human genome. It comprises ten coding genes (*PSG1-9, 11*) and one predominantly non-coding gene (*PSG10*) that are expressed in the placenta and gut, in addition to several poorly characterized long non-coding RNAs. We report that long non-coding RNA *PSG8-AS1* has an oligodendrocyte-specific expression pattern and is co-expressed with genes encoding key myelin constituents. *PSG8-AS1* exhibits two peaks of expression during human brain development coinciding with the most active periods of oligodendrogenesis and myelination. *PSG8-AS1* orthologs were found in the genomes of several primates but significant expression was found only in the human, suggesting a recent evolutionary origin of its proposed role in myelination. Additionally, because co-deletion of chromosomes 1p/19q is a genomic marker of oligodendroglioma, expression of *PSG8-AS1* was examined in these tumors. *PSG8-AS1* may be a promising diagnostic biomarker for glioma, with prognostic value in oligodendroglioma.

## 1 Introduction

Segmental duplications, comprising approximately 7% of the human genome, drive rapid gene evolution and functional innovation in primates, and there is evidence that novel genes within these regions have contributed to the expansion of the neocortex, a feature uniquely developed in the human brain (Bailey and Eichler, 2006; Florio et al., 2018; Fowler, 2022; Samonte and Eichler, 2002; Stankiewicz et al., 2004). While several protein coding genes in segmental duplications have been implicated, modern evolutionary genetic studies indicate that biological innovation is also underpinned by the extensive non-coding regions found in genomes (Jo and Choi, 2015; Mattick, 2001; Mattick et al., 2023). In this context, long non-coding RNAs (lncRNAs) have been described with essential roles in regulating gene expression through diverse mechanisms (Yao et al., 2019). Many lncRNAs are cell and developmental-stage specific and exhibit rapid turnover across species (Mattick et al., 2023). Around 40% of human lncRNAs are expressed in the brain and play crucial roles in neural cell lineage differentiation and function (Aprea et al., 2013; Belgard et al., 2011; Derrien et al., 2012; Necsulea and Kaessmann, 2014; Ninou et al., 2021; Prajapati et al., 2020; Wu et al., 2010). Dysregulated lncRNAs in the human central nervous system (CNS) have been associated with various neurological and neuropsychiatric disorders, underlining the importance of lncRNAs in normal brain development and function (Cai et al., 2020; Faghihi et al., 2008; Jain et al., 2016; Katsel et al., 2019; Katsushima et al., 2016; Lipovich et al., 2012; Talkowski et al., 2012; Voce et al., 2019; Yue et al., 2019).

Despite extensive research in the mouse, which has revealed an essential role for non-coding genes in glial cell development and myelination (Dong et al., 2015; He et al., 2017; Kasuga et al., 2019; Mercer et al., 2010; Stolt et al., 2005; Tochitani and Hayashizaki, 2008; Wei et al., 2021), it is unclear whether lncRNAs act similarly in the human. The scarcity of appropriate models in which to study human-specific lncRNAs means that elucidating their functions in brain development and evolution is challenging (Cabili et al., 2011; Paralkar et al., 2014). In this study, we report the preliminary characterization of lncRNA *PSG8-AS1*, which is named for its complementarity to a minor splice variant of the *PSG8* gene transcript, and not because of any known functional interaction with *PSG* genes. Leveraging publicly available brain gene expression data, including bulk, single-cell, and spatial RNA-seq resources, we identified *PSG8-AS1* as a human oligodendrocyte specific lncRNA potentially implicated in myelination. Additionally, we report the association of *PSG8-AS1* expression with the diagnosis and prognosis of oligodendroglioma, which is genomically characterized by a 1p/19q codeletion together with other genetic markers (Eckel-Passow et al., 2015; The Cancer Genome Atlas Research Network, 2015).

## 2 Methods

### 2.1 Gene expression datasets

Gene expression and exon usage data for *PSGs* and *PSG8-AS1* in adult tissues were accessed through the GTEx Portal (www.gtexportal.org). Expression data used were gene-level transcripts per million (TPM) quantifications generated by the GTEx consortium (Lonsdale et al., 2013). BrainSpan was accessed through the Allen Brain Atlas portal (https://portal.brain-map.org/). Normalized expression data in RPKM (reads per kilobase of exon model per million mapped reads) generated by the BrainSpan resource was used for analyses (Miller et al., 2014). The Cancer Genome Atlas (TCGA, PanCancer Atlas) Low Grade Gliomas (LGG) collection, comprising 514 LGG brain tumor samples subjected to RNA-seq was accessed through TCGAbiolinks R package (The Cancer Genome Atlas Research Network, 2015). Gene counts were normalized by the trimmed mean of M-values (TMM) method (Robinson and Oshlack, 2010). Publicly available datasets that did not belong to a public consortium or atlas were accessed through the Gene Expression Omnibus (GEO) database (https://www.ncbi.nlm.nih.gov/geo/) with accession numbers GSE109082, GSE148241, GSE67835, GSE118257, GSE100796, GSE30352, GSE165595, GSE59612. For a full description of the datasets, see Supplementary Table 1.

### 2.2 Human oligodendrocyte differentiation from induced pluripotent stem cells

Human oligodendrocytes and oligodendrocyte progenitor cells (OPCs) were derived from induced pluripotent stem cell (hiPSC) lines, which were kindly provided by Dr. Elena Gonzalez-Munoz. For experimental details, please see Supplementary Material.

### 2.3 Gene conservation and expression in primates

The online tool blastn (https://blast.ncbi.nlm.nih.gov/Blast.cgi) was used to retrieve potential *PSG8-AS1* orthologs. Sequences (encompassing genomic DNA, cloned cDNAs and predicted RNAs from all available species, i.e, the *nr* collection) aligning to human *PSG8-AS1* RNA sequence (NR_036584.1) were filtered by at least 80% alignment span and similarity. Synthetic constructs were excluded. Status of the genome sequence of *PSG8-AS1* across primate species was explored in the UCSC browser Mammals Multiz Alignment & Conservation (27 primates) track. Multi-Alignment of the RNA sequences of *PSG8-AS1* orthologs was performed with the msa package in R, using the ClustalW algorithm (Thompson et al., 1994). Expression of *PSG8-AS1* orthologs was assessed by the realignment of raw sequencing reads from GSE100796 and GSE30352 (SRP111096 and SRP007412 Sequence Read Archives, respectively) to the genomes of the human (hg38), gorilla (gorGor6), orangutan (ponAbe3), gibbon (nomLeu3), and rhesus macaque (rheMac10) with STAR v2.7 software. In the case of the rhesus macaque, the gene annotation file was modified to incorporate a putative *PSG8-AS1* ortholog. Gene read counts were converted to reads per million (RPM) in R.

### 2.4 Weighted co-expression network analysis and functional annotation

A weighted co-expression network was constructed using the WGCNA package in R (Langfelder and Horvath, 2008). Genes with extremely low expression were removed (genes were kept when the expression was higher than 0.01 RPKM across two thirds of the samples), resulting in 24,267 remaining genes. Soft-thresholding power of 7 was selected. The adjacency matrix was calculated using the Pearson correlation coefficient, and the topological overlap matrix (TOM) was computed to identify highly connected modules. The modules were identified using the dynamic tree-cutting algorithm with a minimum module size of 30 and a merge cut height value of 0.25. A total of 48 modules were identified. The *PSG8-AS1* containing module included a total of 365 genes (Supplementary Table). The functional annotation online tool from the Database for Annotation, Visualization and Integrated Discovery (DAVID) (https://david.ncifcrf.gov/) (Dennis et al., 2003) was used to perform gene-set enrichment analyses.

### 2.5 Glioma expression and survival analyses

Overall expression and mutation data of *PSG8-AS1* in tumors was examined with the cBioPortal online tool (https://www.cbioportal.org/). To evaluate if *PSG8-AS1* was downregulated in glioma, its expression in tumor and non-tumor tissue from glioma patients and controls was compared in datasets GSE165595 and GSE59612. Several glioma subtypes were included to assess the diagnostic value across different molecular subtypes. The Kruskal-Wallis Test (a non-parametric alternative to one-way ANOVA) followed by Pairwise Wilcoxon Rank Sum Tests were performed to evaluate significant differences between the groups. To examine prognosis, clinical data from the TGCA was retrieved with the TCGAbiolinks R package. Samples were kept only if there was information about their gene expression and clinical status. Samples were classified based on their molecular subtype (IDH-wt, IDH-mt alone, and IDH-mt with 1p/19 codeletion) and their expression of *PSG8-AS1*. For each molecular subtype, the samples at the top 25% of *PSG8-AS1* expression were termed “High”, and the bottom 25% were termed “Low”. Kaplan Meier curves and Log-rank tests were then created and visualized with survival and survminer R packages.

## 3 Results

### 3.1 *PSG8-AS1* expression analysis suggests oligodendrocyte specificity

A possible functional relationship between *PSG8-AS1* and the coding genes that constitute the segmental duplication of the *PSG* locus was initially examined by comparing their respective tissue expression patterns. According to GTEx, *PSG8-AS1* is considerably enriched in all compartments of the brain, in contrast to coding *PSGs*, which are highly expressed in the placenta (datasets GSE109082 and GSE148241) and in other tissues at lower levels (GTEx) (Figure 1A-B) (Lonsdale et al., 2013). Antisense lncRNAs are known to regulate the transcription of parental mRNA (Boque-Sastre et al., 2015). However, due to the non-overlapping tissue expression patterns and the extremely low transcription of the *PSG8* intron which *PSG8-AS1* overlaps, downregulation of *PSG8* expression by *PSG8-AS1* seems unlikely (Figure 1A-B) (Supplementary Figure 1).

**Figure 1.**
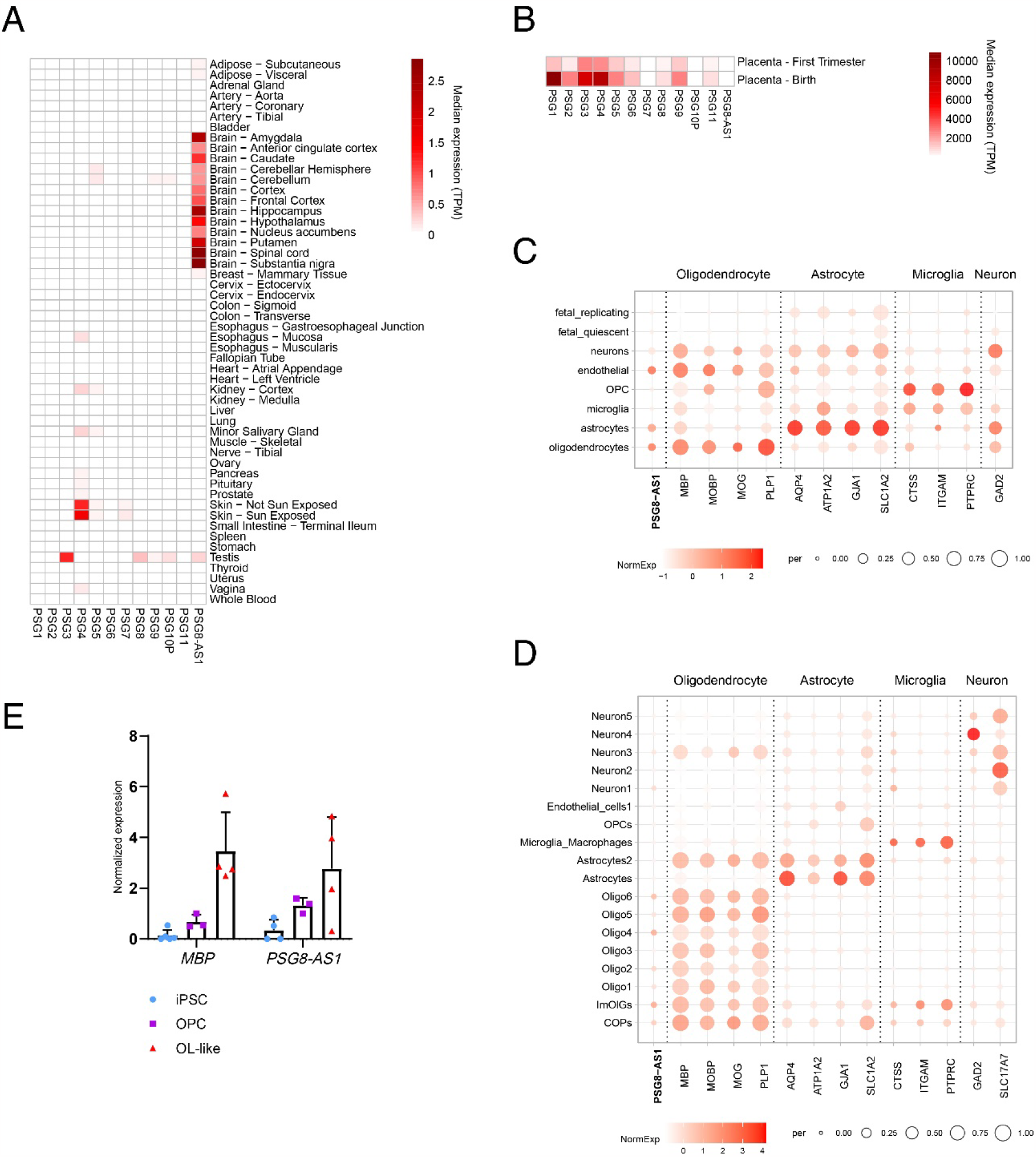
Tissue and cell type-specific expression of *PSG*s and *PSG8-AS1*. A) Heatmap of median expression in Transcripts per Million (TPM) for each gene and tissue from GTEx. B) Heatmap of median expression in TPM for each gene in two placenta datasets (GSE109082 and GSE148241). C-D) Dotplots of scRNA-seq normalized expression of different brain cell markers and *PSG8-AS1* across brain cell types. Expression is shown as colour intensity; percentage of cells where gene expression is identified is represented by size of circle in dotplot. Data from GSE67835 (C) and GSE118257 (D). E) Expression of myelin marker *MBP* and *PSG8-AS1* in human induced pluripotent stem cells (hiPSC), oligodendrocyte progenitor cells (OPC) and oligodendrocytes at late states of differentiation *in vitro* (OL-like), measured by qPCR. Each differentiation stage was represented by at least 3 biological replicates. Data represented as ΔΔCt (individual values, mean +SD), normalised against *GAPDH* and one of the OPC biological replicates.

There is evidence that lncRNAs are cell type-specific (Cabili et al., 2011). Cell specificity was analyzed using two publicly available brain single cell RNA sequencing (scRNA-seq) datasets containing cell identity information (GSE67835 and GSE118257). *PSG8-AS1* was preferentially expressed in oligodendrocytes across all cell subtypes, despite being less abundant than marker genes in both datasets (Figure 1 C-D). This observation can be explained by the fact that lncRNAs are more challenging to detect in scRNA-seq experiments due to their low expression levels compared to coding RNAs (Cabili et al., 2011). When *PSG8-AS1* expression is present in other cell types, such as endothelial cells, oligodendrocyte markers are also expressed (Figure 1C-D). The Brain Cell Type Specific Gene Expression R/Shiny tool (http://celltypes.org/brain/) (McKenzie et al., 2018) also confirms oligodendrocyte specificity (Supplementary Figure 2). Moreover, similar to myelination marker *MBP* (Lopez-Caraballo et al., 2020), we found enriched expression of *PSG8-AS1* in differentiated oligodendrocytes compared to oligodendrocyte progenitor cells (OPCs) and human induced pluripotent stem cells (hiPSCs), confirming our bioinformatic analyses (Figure 1E) (Lopez-Caraballo et al., 2020).

Transcription factor binding to ENCODE cCREs in proximity to the genomic sequence of *PSG8-AS1* was scrutinised to further examine its specificity for oligodendrocytes (Supplementary Table 2). The RE1-Silencing Transcription factor (REST) complex, which represses transcription of genes underpinning neural cell fate decisions in non-neural tissues, and ZNF24 and TCF7L2, two transcription factors essential for terminal oligodendrocyte differentiation and myelination (Abrajano et al., 2009; Elbaz et al., 2018, p. 24; Zhang et al., 2021), bind regulatory regions that are adjacent to brain eQTLs for *PSG8-AS1* (Supplementary Figure 3). There is also evidence for the tumor suppressor protein p53 binding the *PSG8-AS1* promoter (Supplementary Figure 3; Supplementary Table 3).

### 3.2 *PSG8-AS1* sequences evolved in the Catarrhini lineage but expression may be confined to the human

Predicted orthologs of *PSG8-AS1* are found in the most recent genome assemblies of the gorilla (Homininae), orangutan (Hominidae), gibbon (Hominoidea), Tibetan macaque, rhesus macaque, drill and baboon (Catarrhini) (Figure 2A). In the chimp and bonobo, our closest primate relatives, *PSG8-AS1* exons 2 and 3 are either not conserved or are poorly assembled (Suplementary Figure 4). While assembly gaps are common in segmental duplications (Samonte and Eichler, 2002; Vollger et al., 2022), it is also possible that the common ancestor between the human, chimp and bonobo had a functional copy of *PSG8-AS1*, which was only retained in the human lineage (Suplementary Figure 4). Conservation at the RNA level was also inspected by multiple alignment of RNA sequences of *PSG8-AS1* orthologs (Figure 2A). This showed that exon sequence composition is not the same across orthologous lncRNAs, with the macaques, drill and baboon losing exon 1 almost entirely, and gibbons losing most of exon 4. To determine if *PSG8-AS1* orthologs are expressed in primates other than the human, RNA-seq reads from brain and other tissues (heart, liver, kidney, testis) from several primate species were aligned to their respective genomes (SRP111096 and SRP007412 Sequence Read Archives, from GSE100796 and GSE30352, respectively). The analysis showed predominant expression of *PSG8-AS1* in the human brain, with residual levels in the brain of non-human primates (Figure 2B). Levels in all tissues in non-human primates were low, similar to non-brain tissues in the human (Figure 2C) This suggests that expression of *PSG8-AS1* orthologs in non-human primates might be negligible compared to the human.

**Figure 2.**
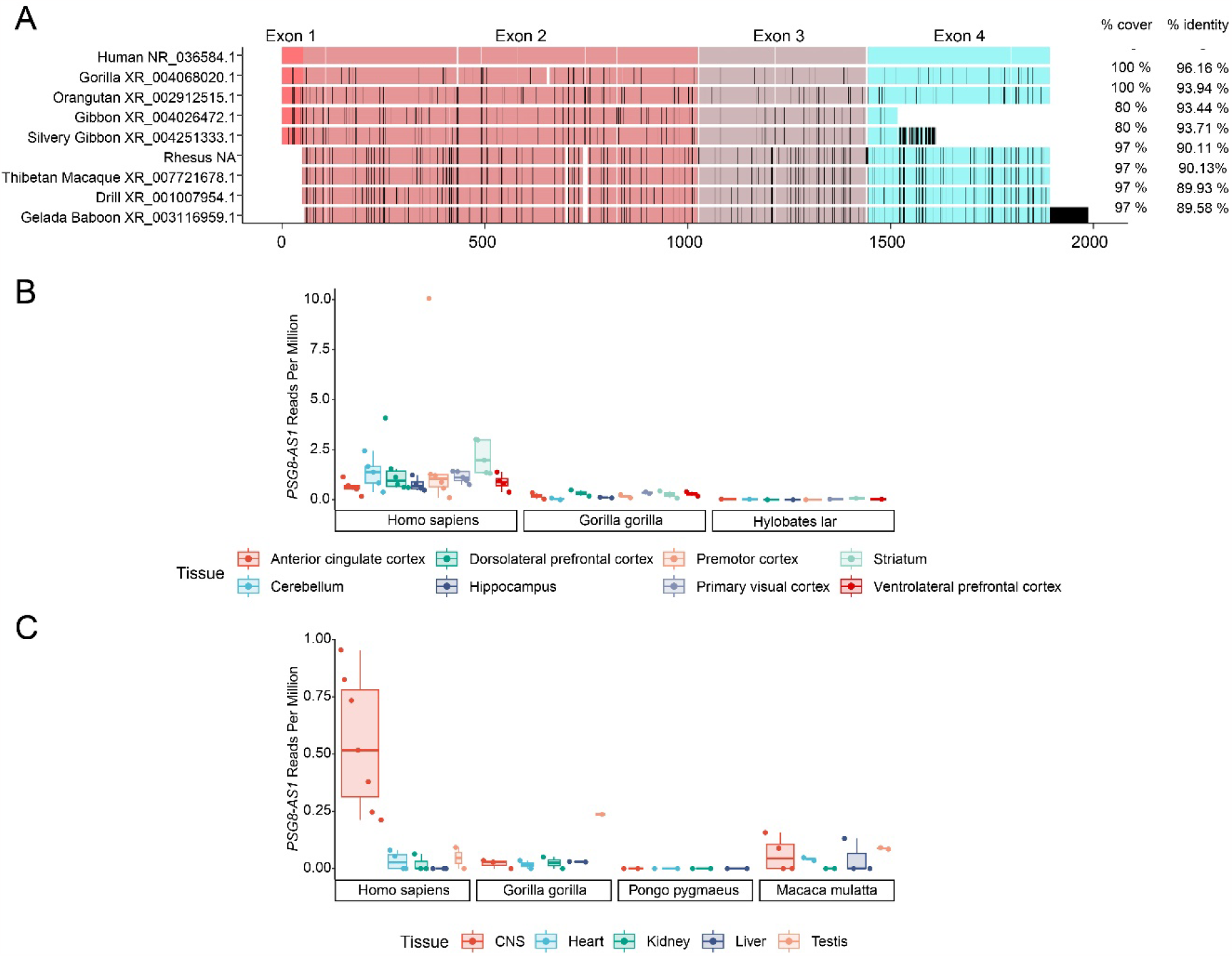
Conservation of exon sequences and RNA expression of *PSG8-AS1* in primates. A) Multi-Alignment of RNA sequences of blastn-predicted *PSG8-AS1* orthologs. % cover and % identity of alignment with human *PSG8-*AS1 shown on the right. Gaps (white), nucleotide substitutions or insertions compared to the human (black); matches in exons (as annotated in the human) colour coded: Exon 1 (pink), Exon 2 (light pink), Exon 3 (purple), Exon 4 (cyan). B-C) Boxplots with individual values showing *PSG8-AS1* and ortholog expression levels in different primate species and tissues. Expression units is Reads per Million (RPM). B: Expression data from different brain regions in human, gorilla, and gibbon (GSE30352). C: Expression data from different tissues, including the central nervous system (CNS), in human, gorilla, orangutan and macaque (GSE100796).

### 3.3 *PSG8-AS1* is co-expressed with myelin constituents in the developing and adult brain

Since many novel genes important for human brain evolution are expressed during development, *PSG8-AS1* expression was examined in the Allen Brain Atlas human brain development transcriptomic resource (Hawrylycz et al., 2012). This dataset includes 31 time points pre- and post-birth from 26 brain regions. Levels of *PSG8-AS1* transcription were compared with specific brain cell markers (McKenzie et al., 2018) (Figure 3A). Coincident with the start of oligodendrogenesis and the period of most rapid myelination in humans (25-27 weeks post-conception [pcw] and 2 years after birth, respectively), *PSG8-AS1* expression peaks at the same time as oligodendrocyte markers *MBP* and *MOG* (de Faria et al., 2021; Ouyang et al., 2019). In contrast, astrocyte markers *GJA2* and *AQP4* start increasing before the initiation of gliogenesis at approximately 24 pcw, similar to neural markers *GAD2* and *SLC17A7* (Figure 3A) (McKenzie et al., 2018).

**Figure 3.**
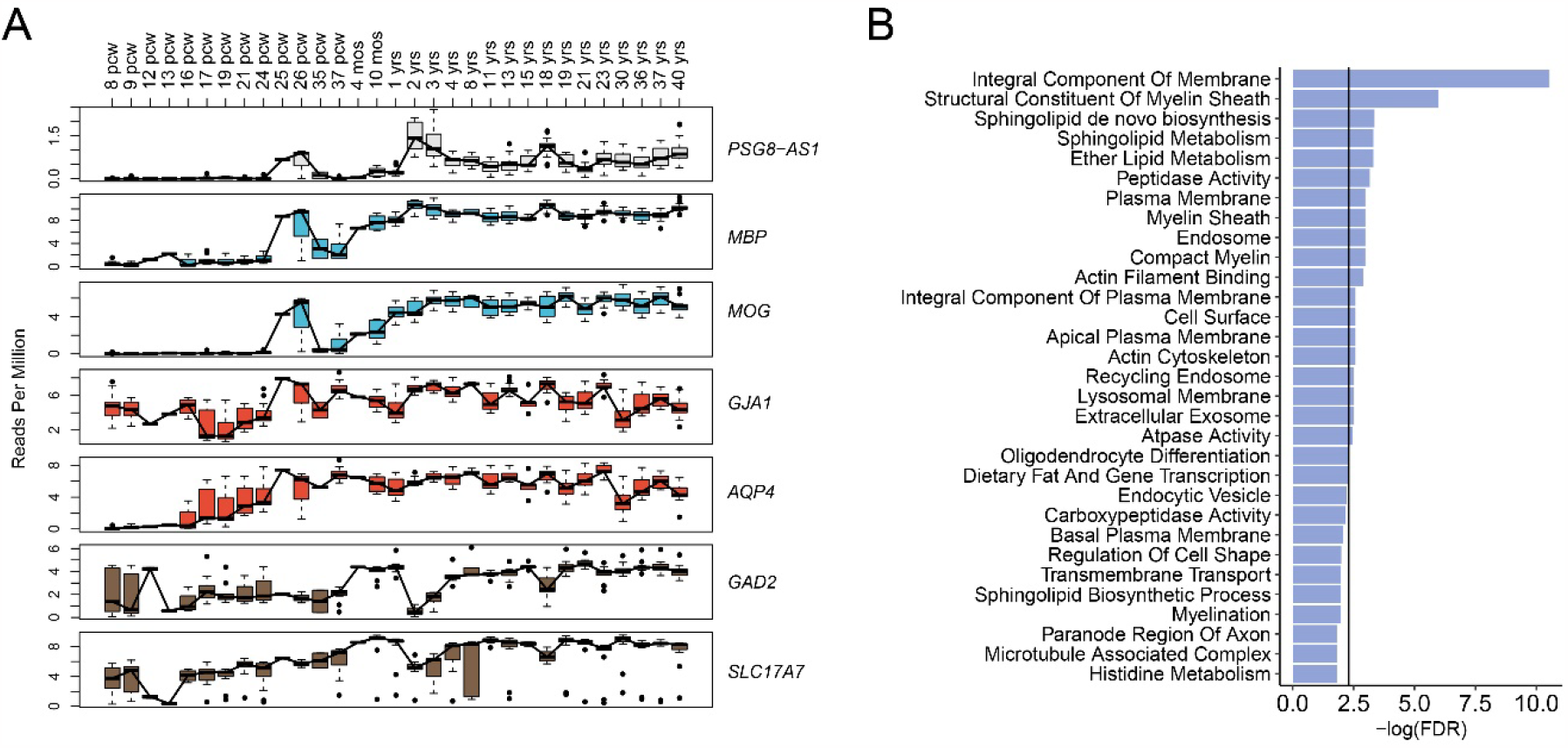
*PSG8-AS1* expression coincides with oligodendrocyte markers and myelin constituents during human brain development. A) Expression of *PSG8-AS1* and several brain cell markers across developmental stages. Expression units are Reads per Million (RPM). *PSG8-AS1* boxplots are shown in grey, oligodendrocyte markers (*MBP* and *MOG*) in blue, astrocyte markers (*GJA1* and *AQP4*) in red, and neuron markers (*GAD2* and *SLC17A7*) in brown. B) Barplot showing False Discovery Rate (FDR) of enriched pathways in the gene set enrichment analysis of the *PSG8-AS1* high-correlated gene expression module. Pathway name format was simplified for visualization (see Supplementary Table 5 for full names). Gene set enrichment analysis was performed with DAVID online tool. Vertical black line represents FDR = 0.1.

To further clarify which genes are co-expressed with *PSG8-AS1* across brain developmental stages, Weighted Gene Correlation Network Analysis (WGCNA) (Langfelder and Horvath, 2008) on the Allen Brain Atlas dataset was performed. The genes most correlated with *PSG8-AS1* within the module were *MBP, FOXO4* and *KIF1C* (Supplementary Table 4). A pathway enrichment analysis was conducted on the *PSG8-AS1* module using the Database for Annotation, Visualization and Integrated Discovery (DAVID) tool (Dennis et al., 2003). The most significant pathways were related to myelination (Structural Constituent of Myelin Sheath, Myelin Sheath, Compact Myelin) and cellular membrane composition (Integral Component of Membrane, Integral Component of Plasma Membrane, Plasma Membrane, Sphingolipid de novo Biosynthesis, Sphingolipid Metabolism) (Figure 3B; Supplementary Table 5). Of note, sphingolipids are crucial for the synthesis and maintenance of the myelin sheath (Giussani et al., 2021). Although only marginally significant, Oligodendrocyte Differentiation was also enriched.

### 3.4 *PSG8-AS1* expression is a potential biomarker in glioma

The location of *PSG8-AS1* on chromosome 19q13 and its association with oligodendrocyte function prompted an examination of its expression in gliomas, and particularly oligodendroglioma, which have been characterized by mutations in *IDH1-2* with 1p/19q codeletions (Eckel-Passow et al., 2015; The Cancer Genome Atlas Research Network, 2015). No germline or somatic mutations were found in *PSG8-AS1* in grade II and III oligodendroglioma and astrocytoma (low-grade gliomas or LGG) by assessing The Cancer Genome Atlas (TCGA) data through cBioPortal (Cerami et al., 2012; Gao et al., 2013; The Cancer Genome Atlas Research Network, 2015). The RNA levels of *PSG8-AS1* in non-tumor and tumor tissue were assessed in two datasets (GSE165595, GSE59612). *PSG8-AS1* expression was lower in any subtype of glioma when compared with paired non-tumor brain tissue (Pairwise Wilcoxon Rank Sum Tests; “non-tumoral” vs. glioblastoma p=0.0037; “non-tumoral” vs. IDHmt p=0.0002; “non-tumoral” vs. IDH-mt 1p/19q codeletion p= 7.6e-05) (Figure 4A). Contrast enhancement on magnetic resonance (MR) images, which is associated with the destruction of the blood-brain barrier and higher malignancy, is specific to high-grade gliomas (Gill et al., 2014; Pronin et al., 1997). When comparing non-neoplastic brain tissue with contrast-enhancing glioma and non-enhancing glioma, the expression of *PSG8-AS1* was only significantly lower in contrast-enhancing samples, suggesting that *PSG8-AS1* could be an indicator of glioma malignancy (Pairwise Wilcoxon Rank Sum Tests; non-neoplastic vs. contrast-enhancing glioma p=6.5e-08; non-neoplastic vs. non-enhancing glioma p=0.13; non-enhancing glioma vs contrast-enhancing glioma p=4.5e-09) (Figure 4B).

**Figure 4.**
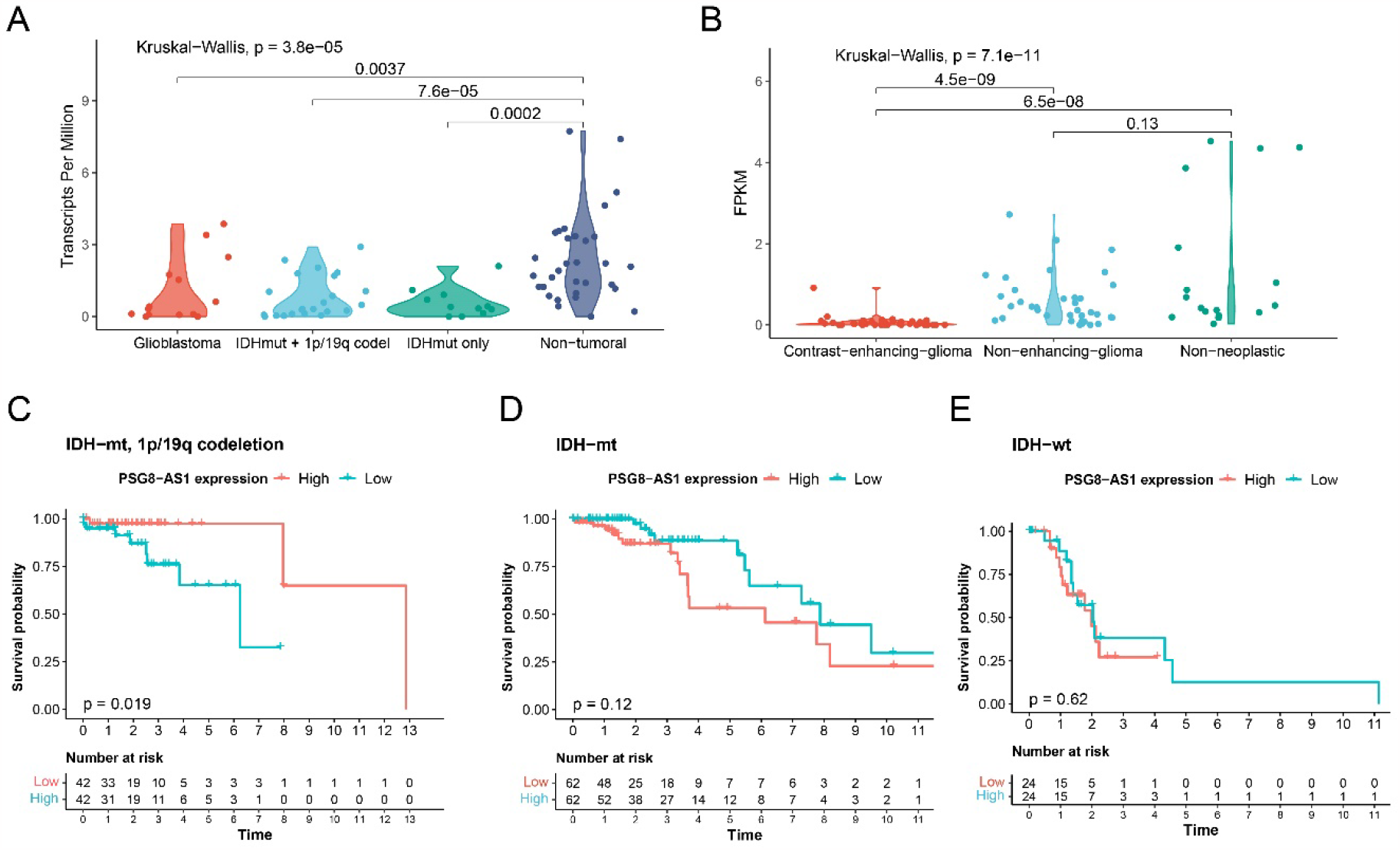
Relevance of *PSG8-AS1* expression in low-grade gliomas. A-B) *PSG8-AS1* expression in tumours compared to paired non-tumoral tissue. Violin plots showing Transcript Per Million (TPM) values from GSE165595 (A) and GSE59612 (B). Individual values are shown in dots. Significant differences between the groups were assessed with a Kruskal-Wallis test (nonparametric one-way ANOVA). C-E) Kaplan-Meier survival curves in patients from the LGG TCGA study stratified by *PSG8-AS1* expression in IDH-mutant, 1p/19q codeleted glioma (C), IDH-mutant glioma without 1/p19q codeletion (D) and IDH-wildtype glioma (E). Time is expressed in years.

To further investigate the potential clinical significance of *PSG8-AS1* downregulation, the association between *PSG8-AS1* expression and survival was examined. TCGA patients were stratified by molecular subtype and *PSG8-AS1* expression (high = top 25% and low = bottom 25% in each molecular subtype). Kaplan-Meier analysis and Log-rank tests were carried out to evaluate the effect on overall survival. In IDH-mt 1p/19q codeleted glioma (oligodendroglioma), high expression levels of *PSG8-AS1* significantly correlated with longer patient survival time (Log-rank p = 0.019) (Figure 4C), whereas there was no effect on survival for the subtypes without the codeletion genotype (Figure 4D-E). The results of these analyses suggest that *PSG8-AS1* levels might have diagnostic value in gliomas and could serve as a prognostic biomarker in oligodendrogliomas.

## 4 Discussion

We characterized *PSG8-AS1*, a lncRNA located in the rapidly evolving *PSG* segmental duplication. *PSG8-AS1* exhibits a non-overlapping expression pattern with the coding genes in the *PSG* locus and is predominantly expressed in the brain. *PSG8-AS1* showed predominant expression in differentiated oligodendrocytes as determined by analysis of brain scRNA-seq data and qPCR on hiPSCs-derived oligodendrocytes. Public databases report binding of components of REST and CoREST and transcription factors ZNF24 and TCF7L2, which are important regulators of oligodendrocyte differentiation and myelination in the human, to putative enhancers of *PSG8-AS1* (Abrajano et al., 2009; Elbaz et al., 2018; Zhang et al., 2021). These observations support the contention that *PSG8-AS1* is an oligodendrocyte-specific gene.

*PSG8-AS1* was hypothesised to be human-specific due to its localization to a segmental duplication, many of which are known to give rise to novel genes important in human brain evolution (Florio et al., 2018; Vollger et al., 2022). Conservation analyses indicated that *PSG8-AS1* may have emerged in the Catarrhini lineage, although its expression is primarily observed in the human. The emergence of genes important for brain evolution in primate genomes with preferential expression in the human brain has been described previously, and it has been speculated that specific changes in regulatory elements occurred in these genes during human evolution (Florio et al., 2018). Additionally, the virtually exclusive focus on neurons when considering recent human brain evolution is now thought to be insufficient to explain human cognitive capabilities (Berto et al., 2019; Donahue et al., 2018; Rilling and van den Heuvel, 2018; Sousa et al., 2017). According to a recent study, oligodendrocytes exhibit more rapid evolution than neurons (Berto et al., 2019), and the human brain exhibits enhanced connectivity compared to non-human primates, which requires a longer period of myelination (Donahue et al., 2018; Miller et al., 2012; Rilling and van den Heuvel, 2018). This, together with the growing evidence linking oligodendrocyte function and myelination to cognition and neuropsychiatric disorders (Fields et al., 2014; Maas et al., 2020; Mighdoll et al., 2015; Voineskos et al., 2013), suggests that the characterization of human oligodendrocyte-specific genes such as *PSG8-AS1* will contribute to our understanding of brain evolution, development and disease. In the developing human brain, *g*enes highly correlated with *PSG8-AS1* were implicated in myelination and the composition of the plasma membrane. The molecular mechanisms by which *PSG8-AS1* might regulate this process remains unclear. In future research, the use of oligodendrocyte progenitors derived from hiPSCs or human cerebral organoids, together with the perturbation of *PSG8-AS1* expression should be considered, as these approaches have been used successfully in previous studies (Lopez-Caraballo et al., 2020; Mahmoud et al., 2019; Pantoja et al., 2020; Shaker et al., 2021; Xu et al., 2021).

We hypothesised that *PSG8-AS1* might be involved in glioma because the tumor suppressor p53 binds a proximal enhancer. In addition, codeletion of 19q, where *PSG8-AS1* is located, together with 1p, is used as a genetic marker for oligodendroglioma (The Cancer Genome Atlas Research Network, 2015). Accordingly, we found that *PSG8-AS1* expression exhibited diagnostic value for all subtypes of glioma, and prognostic value specifically for oligodendroglioma. The effect of *PSG8-AS1* expression on overall survival of oligodendroglioma, but not other gliomas, is consistent with a role for *PSG8-AS1* in oligodendrocyte function and implies that *PSG8-AS1* levels are reduced in oligodendroglial tumor cells with increased malignancy. It has been suggested that oligodendroglioma-specific tumor suppressor genes map to chromosomes 1p/19q (Bou Zerdan and Assi, 2021). For example, recurrent mutation of the *FUBP1* and *CIC* genes, located on chromosomes 1p and 19q respectively, have been found in oligodendrogliomas and linked to more aggressive forms of the tumor (Bettegowda et al., 2011; Gladitz et al., 2018). Of note, because of karyotypic differences between the human and other species, this genomic signature of oligodendrogliomas is most likely unique to the human (Lv et al., 2021).

In summary, this study provides the first in-silico characterization of *PSG8-AS1*, a human oligodendrocyte-specific lncRNA mapping to the *PSG* segmental duplication on chromosome 19q and suggests a role in myelination. Our findings also suggest potential biomarker utility for *PSG8-AS1* in oligodendrogliomas. Further study will be required to elucidate the precise mechanisms by which *PSG8-AS1* regulates myelination and to explore its potential as a diagnostic and therapeutic target in myelination disorders and cancer.

## 6 Author Contributions

MBR and TM contributed to conceptualization of the study and writing the manuscript. MBR performed bioinformatic analyses and qRT-PCRs. EGM performed hiPSC differentiation and provided RNA samples.

