## Supplemental Material for "Oligodendrocyte-specific expression of *PSG8-AS1* suggests a role in myelination with prognostic value in oligodendroglioma"

Supplementary Material

### 1 Supplementary Methods

#### 1.1 Human induced pluripotent stem cell differentiation

All cell lines were extensively validated in previous studies (Douvaras and Fossati, 2015; Lopez-Caraballo et al., 2020). Differentiation of hiPSCs into OLs was carried out according to the “original protocol” published by (Douvaras and Fossati, 2015) with slight modifications. Briefly, undifferentiated iPSCs were cultured in mTeSR1 cell medium (StemCell Technologies, cat. no. 05850). iPS cell colonies were then dissociated into single cells using Accutase (Life Technologies, Carlsbad, CA, United States, #A11105-01), seeded in matrigel coated 6-well plates at 2.5 x 10 ^5^ cells/well density, and cultured in mTeSR1 medium for 2 days before differentiation, which was induced by switching to neural induction medium (NIM) containing freshly added SB434542 (TGFb inhibitor), LDN-193189 (BMP inhibitor), and retinoic acid (RA) (day 0). Medium was changed daily for 8 days. From day 8 to 12, cells were exposed to RA and smoothened agonist (SAG). On day 12, overconfluent cells forming 3D structures were clearly visible, and the cells were mechanically dissociated, transferred to non-coated 6-well plates and grown in suspension until day 20 in N2B27 medium with 25 μg/ml insulin, 100 nM RA and 1 μM SAG. On day 20, cells were switched to PDGF containing medium until day 30 (OPC stage). On day 30, cell aggregates were replated onto poly-L-ornithine- and laminin-coated 6-well plates and grown adherent through day 75. On Day 76, cells were switched to mitogen-free medium for the rest of the differentiation (OL stage) and further used for RNA extraction and qPCR analysis.

#### 1.2 RNA isolation and quantitative Polymerase Chain Reaction (qPCR)

RNA isolation was performed with RNeasy kit (Qiagen) following manufacturer’s instructions. For cDNA synthesis, High-Capacity cDNA Reverse Transcription Kit (Applied Biosystems™) was used following manufacturer’s protocol. Gene expression was then quantified by qPCR with the ChamQ Universal SYBR qPCR Master Mix system (Vazyme) using primers listed in Supplementary Table 6. Data analysis was performed by applying the ΔΔct method, normalizing against house-keeping gene *GAPDH* and one of the OPC biological replicates. For each differentiation stage (iPSCs, OPCs and oligodendrocytes) at least three biological replicates were used.

#### 1.3 Transcription factor binding and expression quantitative trait loci (eQTLs)

The University of California, Santa Cruz Genome Browser (UCSC Browser, <https://genome.ucsc.edu/>) was used to visualize the genomic location of regulatory regions and eQTLs potentially influencing *PSG8-AS1* expression in hg38. Candidate cis-regulatory elements (cCREs), which include promoters, enhancers and CTCF-binding elements were retrieved from the ENCODE v3 library (Moore et al., 2020). Transcription factor binding information from the same ENCODE library was integrated with the ORegAnno database (Lesurf et al., 2016). Transcription factors were classified by their summary in the GeneCards database (<https://www.genecards.org/>). If the central nervous system was mentioned, the protein was classified as CNS regulation. If the REST complex was mentioned, the protein was classified as REST complex. eQTLs were obtained from GTEx and PhychENCODE resources (Akbarian et al., 2015; Lonsdale et al., 2013).

### 2 Supplementary Data

#### 2.1 Supplementary Figures

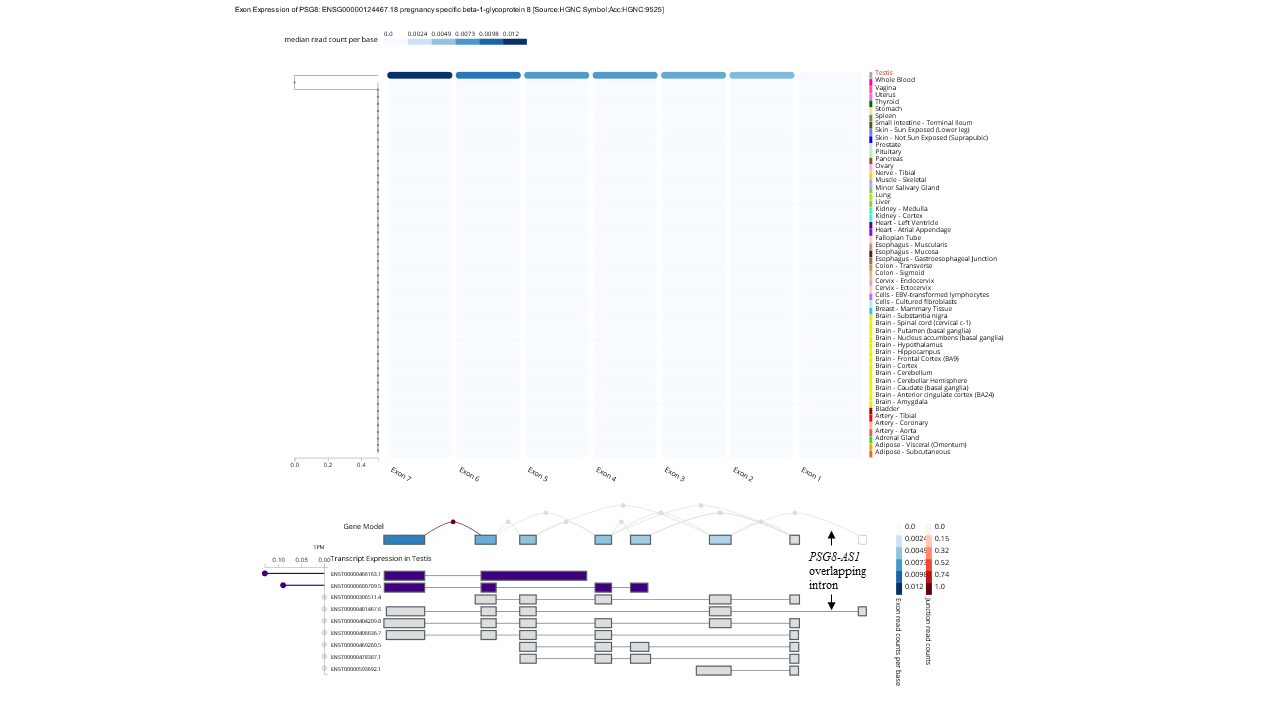

**Supplementary Figure 1.** Exon expression of *PSG8* across tissues. The exon that involves *PSG8-AS1* overlapping intron is never detected in GTEx data (Lonsdale et al., 2013).

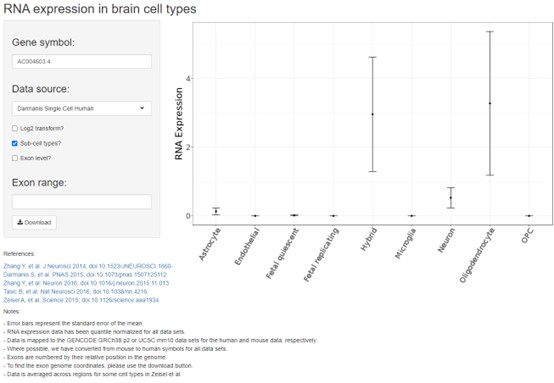

**Supplementary Figure 2.** Validation of *PSG8-AS1* expression specificity in oligodendrocytes. The Brain Cell Type Specific Gene Expression R/Shiny tool was used (http://celltypes.org/brain/) (McKenzie et al., 2018). AC004603.4 is a previous gene name for *PSG8-AS1*.

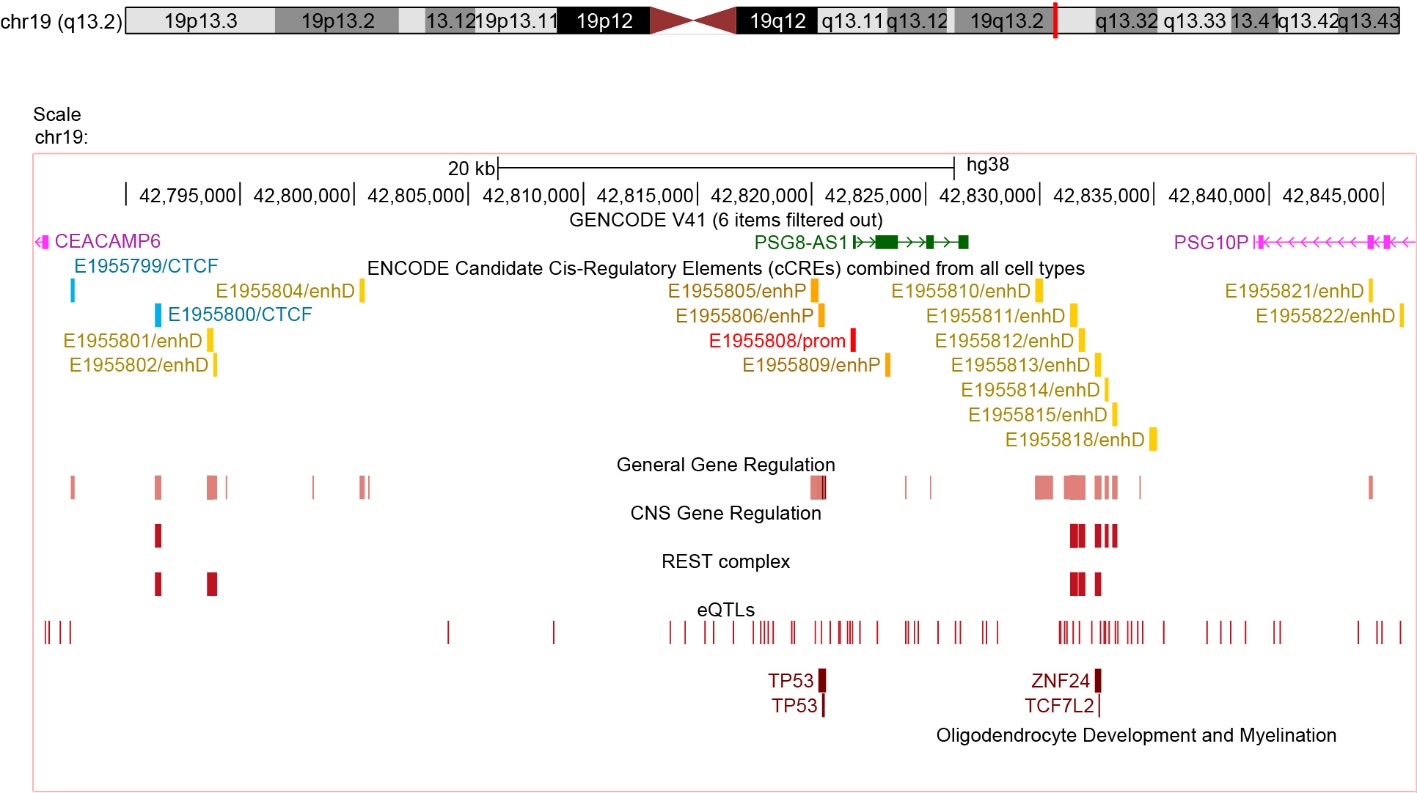

**Supplementary Figure 3.** Localization of cCREs, transcription factor binding sites, and eQTLs that potentially regulate *PSG8-AS1*. Modified from UCSC browser (genome version hg38). GENCODE V41 track shows lncRNA genes in green and pseudogenes in pink. ENCODE cCREs track shows CTCF-bound (blue), distal enhancers (light yellow), proximal enhancers (orange), and promoters (red). Transcription factor binding sites (General, CNS, REST complex, TP53, ZNF24 and TCF7L2) and eQTLs are shown in different shades of red.

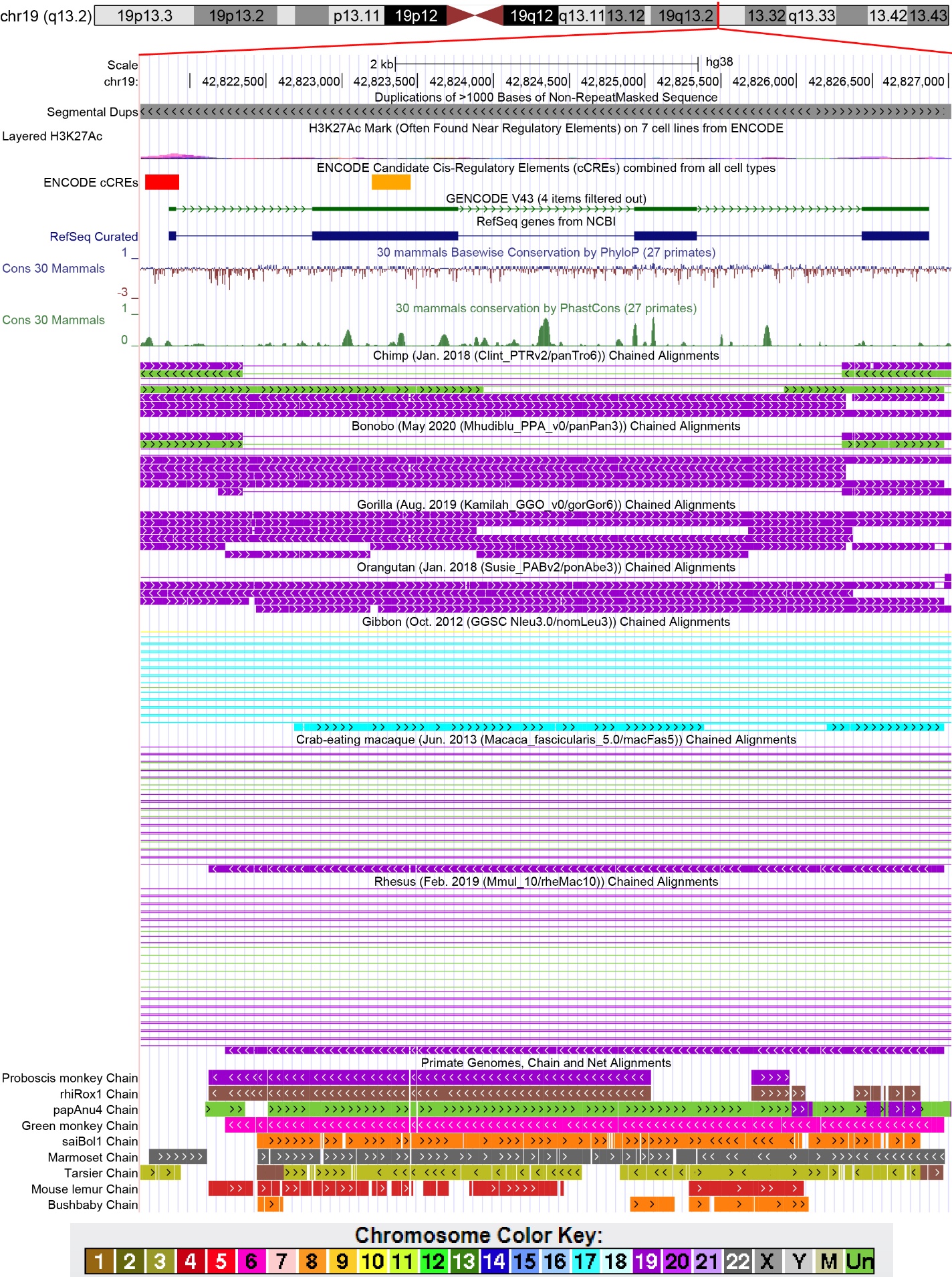

**Supplementary Figure 4.** Alignments of several primate genomes to the human genome (hg38) in the *PSG8-AS1* region in the UCSC browser Mammals Multiz Alignment & Conservation (27 primates). Briefly, a Large-Scale Genome Alignment Tool (lastz) was used to generate pairwise alignments with the human genome (Kent, 2002). Pairwise alignments are then organized into gapless “chains” and scored by a Nearest Neighbour Algorithm (Chiaromonte et al., 2002). Alignment “nets” show the highest-scoring chain at the top, filling the gaps with lower-scoring chains (Kent, 2002). For chimp, bonobo, gorilla, orangutan, gibbon and macaque species all “chains” are shown. For the rest, only “nets” are shown. *PSG8-AS1* sequence and promoter in the human genome is shown at the top (boxes=exons, lines=introns, arrows=strand). For alignments, boxes represent ungapped alignments; lines represent gaps. In the chimp, bonobo, gorilla, orangutan and macaques, *PSG8-AS1*-like sequences reside mostly on chromosome 19. Arrows indicate the direction of the alignment.

#### 2.2 Supplementary Tables (see attached excel file)

**Supplementary Table 1**. Gene expression datasets accessed through GEO.

**Supplementary Table 2**. Transcription factor (TF) binding to cCREs in proximity to *PSG8-AS1* transcription start site (TSS).

**Supplementary Table 3**. Cis-eQTLs for *PSG8-AS1* in the brain (hg38).

**Supplementary Table 4**. Correlation with *PSG8-AS1* of genes in *PSG8-AS1* WGCNA module.

**Supplementary Table 5**. Gene set enrichment analysis performed with DAVID online tool of the *PSG8-AS1* WGNCA high-correlated gene expression module.

**Supplementary Table 6.** Primers used for qPCRs.

Lonsdale, J., Thomas, J., Salvatore, M., Phillips, R., Lo, E., Shad, S., Hasz, R., Walters, G., Garcia, F., Young, N., Foster, B., Moser, M., Karasik, E., Gillard, B., Ramsey, K., Sullivan, S., Bridge, J., Magazine, H., Syron, J., Fleming, J., Siminoff, L., Traino, H., Mosavel, M., Barker, L., Jewell, S., Rohrer, D., Maxim, D., Filkins, D., Harbach, P., Cortadillo, E., Berghuis, B., Turner, L., Hudson, E., Feenstra, K., Sobin, L., Robb, J., Branton, P., Korzeniewski, G., Shive, C., Tabor, D., Qi, L., Groch, K., Nampally, S., Buia, S., Zimmerman, A., Smith, A., Burges, R., Robinson, K., Valentino, K., Bradbury, D., Cosentino, M., Diaz-Mayoral, N., Kennedy, M., Engel, T., Williams, P., Erickson, K., Ardlie, K., Winckler, W., Getz, G., DeLuca, D., MacArthur, D., Kellis, M., Thomson, A., Young, T., Gelfand, E., Donovan, M., Meng, Y., Grant, G., Mash, D., Marcus, Y., Basile, M., Liu, J., Zhu, J., Tu, Z., Cox, N.J., Nicolae, D.L., Gamazon, E.R., Im, H.K., Konkashbaev, A., Pritchard, J., Stevens, M., Flutre, T., Wen, X., Dermitzakis, E.T., Lappalainen, T., Guigo, R., Monlong, J., Sammeth, M., Koller, D., Battle, A., Mostafavi, S., McCarthy, M., Rivas, M., Maller, J., Rusyn, I., Nobel, A., Wright, F., Shabalin, A., Feolo, M., Sharopova, N., Sturcke, A., Paschal, J., Anderson, J.M., Wilder, E.L., Derr, L.K., Green, E.D., Struewing, J.P., Temple, G., Volpi, S., Boyer, J.T., Thomson, E.J., Guyer, M.S., Ng, C., Abdallah, A., Colantuoni, D., Insel, T.R., Koester, S.E., Little, A.R., Bender, P.K., Lehner, T., Yao, Y., Compton, C.C., Vaught, J.B., Sawyer, S., Lockhart, N.C., Demchok, J., Moore, H.F., 2013. The Genotype-Tissue Expression (GTEx) project. Nat. Genet. 45, 580–585. https://doi.org/10.1038/ng.2653

Lopez-Caraballo, L., Martorell-Marugan, J., Carmona-Sáez, P., Gonzalez-Munoz, E., 2020. iPS-Derived Early Oligodendrocyte Progenitor Cells from SPMS Patients Reveal Deficient In Vitro Cell Migration Stimulation. Cells 9, 1803. https://doi.org/10.3390/cells9081803

McKenzie, A.T., Wang, M., Hauberg, M.E., Fullard, J.F., Kozlenkov, A., Keenan, A., Hurd, Y.L., Dracheva, S., Casaccia, P., Roussos, P., Zhang, B., 2018. Brain Cell Type Specific Gene Expression and Co-expression Network Architectures. Sci. Rep. 8, 8868. https://doi.org/10.1038/s41598-018-27293-5

Moore, J.E., Purcaro, M.J., Pratt, H.E., Epstein, C.B., Shoresh, N., Adrian, J., Kawli, T., Davis, C.A., Dobin, A., Kaul, R., Halow, J., Van Nostrand, E.L., Freese, P., Gorkin, D.U., Shen, Y., He, Y., Mackiewicz, M., Pauli-Behn, F., Williams, B.A., Mortazavi, A., Keller, C.A., Zhang, X.-O., Elhajjajy, S.I., Huey, J., Dickel, D.E., Snetkova, V., Wei, X., Wang, X., Rivera-Mulia, J.C., Rozowsky, J., Zhang, Jing, Chhetri, S.B., Zhang, Jialing, Victorsen, A., White, K.P., Visel, A., Yeo, G.W., Burge, C.B., Lécuyer, E., Gilbert, D.M., Dekker, J., Rinn, J., Mendenhall, E.M., Ecker, J.R., Kellis, M., Klein, R.J., Noble, W.S., Kundaje, A., Guigó, R., Farnham, P.J., Cherry, J.M., Myers, R.M., Ren, B., Graveley, B.R., Gerstein, M.B., Pennacchio, L.A., Snyder, M.P., Bernstein, B.E., Wold, B., Hardison, R.C., Gingeras, T.R., Stamatoyannopoulos, J.A., Weng, Z., 2020. Expanded encyclopaedias of DNA elements in the human and mouse genomes. Nature 583, 699–710. https://doi.org/10.1038/s41586-020-2493-4
